# Caffeic acid promoted deep vein thrombosis resolution in mice by suppressed macrophage M1 polarization through Keap1/Nrf2 pathway

**DOI:** 10.1101/2025.07.24.666694

**Authors:** Zhou-Yu Nie, Jia-Qi Zhang, Jia-Yi Shen-Yuan, Meng-Jiao Lu, Yang Shen, Liang Shi, Yong-Bing Cao, Li-Chao Zhang, Ling Li

## Abstract

**Background:** Deep vein thrombosis (DVT) carries significant health risks, with macrophages playing a key role in thrombus resolution. Caffeic acid (CA) has been shown to inhibit thrombosis, but its role in DVT resolution remains unclear.

**Purpose:** This study aimed to investigates the effects and mechanisms of CA in accelerating DVT resolution.

**Methods:** A stasis-type DVT model was established in mice via inferior vena cava ligation. The effects of CA were analyzed. Thrombus resolution was assessed using histological staining, gelatin zymography, immunoblotting, and immunofluorescence. Macrophage depletion was conducted with chlorophosphate liposomes, and bone marrow-derived macrophages (BMDMs) were used for polarization studies. RNA sequencing identified potential molecular pathways. Nrf2 gene-deficient mice were used to verify the role of Nrf2 in deep vein thrombus resolution.

**Results:** Caffeic acid reduced thrombus size, enhanced collagenolysis via increased MMP-2 activity, promoted neovascularization, increased macrophage infiltration, and suppressed M1 polarization and inflammation. CA activated the nuclear factor erythroid 2-related factor 2 (Nrf2) pathway by inhibiting Keap1, leading to enhanced antioxidant responses in both BMDMs and thrombus tissue. Macrophage depletion negated CA’s benefits, confirming the central role macrophages.

**Conclusion:** CA promoted early thrombus resolution via macrophage recruitment, M1 polarization suppression, and Nrf2 activation, highlighting its potential as a preventive strategy for DVT.

## Introduction

Deep vei<u>n</u> thrombosis (DVT) is a frequent, multifactorial disease and a leading cause of morbidity and mortality.^1^ The incidence of DVT in the general population is estimated to be 67 per 100,000 person-years,^2^ and the incidence of venous thromboembolism increases with age.^3^ Current treatments for DVT include anticoagulation therapy, thrombolytic therapy, surgical thrombectomy, inferior vena cava (IVC) filter placement, and others.^4^ However, these treatments have some drawbacks, such as increasing the risk of bleeding, being limited to fresh thrombus, not eliminating the occurrence of post thrombotic syndrome, an absence of acceleration in thrombus resolution, etc.^5^ Therefore, there is an urgent need to develops strategies that directly promote thrombus resolution.

Current researches have found that promoting the natural resolution of formed thrombus is a new strategy for the treatment of DVT.^6^ Monocytes/macrophages in the blood play a critical role in thrombus resolution.^7,8^ It was shown that along with venous thrombosis, the number of monocytes/macrophages recruited into the thrombus is gradually increasing, and the increase in their number is positively correlated with the rate of thrombus resolution.^9,10^ Monocytes/macrophages promote thrombus resolution mainly by phagocytosis of necrotic tissues or cellular debris, secretion of a variety of inflammatory factors, growth factors, and protein hydrolyzing enzymes, which elicit inflammatory responses and promote vascular neogenesis, fibrin and extracellular matrix dissolution.^11,12^ However, an excessive pro-inflammatory response inhibits thrombus resolution.^13^ Therefore, finding the potential candidate drugs that could enhance monocyte/macrophage infiltration into thrombus to accelerate thrombus resolution may possess important clinical implications.

Caffeic acid (CA), [(E)-3-(3, 4-dihydroxyphenyl) propyl-2-enoic acid], a hydroxycinnamic acid whose molecular weight is 180.16 g/mol.^14^ Caffeic acid is the main small molecule compound in the human diet and is widely present in coffee, blueberries, kiwifruit, plum-fruits, cherryfruits and apples.^15–17^ CA is also found in cereals, carrots, salads, eggplant, cabbage and artichokes. Moreover, CA is a natural herbal monomer extracted from Chinese herbs such as *Salvia miltiorrhiza* and propolis, which has been used in traditional Chinese medicine for centuries.^18,19^ Numerous studies have shown that CA has amazing biological activities such as anti-inflammatory, antioxidant, antibacterial, antiviral and anti-cancer.^20–22^ CA has been demonstrated to exert inhibitory effects on thrombus formation in murine cerebral arteries induced by photochemical injury, as well as on cerebral venular thrombosis triggered by topical ADP application.^23^ Furthermore, in vitro experiments revealed its capacity to suppress lipopolysaccharide (LPS)-stimulated M1 polarization in RAW264.7 macrophages.^24^ However, the role of CA in venous thrombus resolution is still unclear.

This study aimed to investigate the ability of CA on deep vein thrombosis resolution and elucidate its underlying mechanisms. Notably, our findings demonstrate that CA significantly enhanced venous thrombus resolution, whereas therapeutic administration of CA post-thrombosis onset exhibited no discernible promotive effect on thrombus resolution. Furthermore, we systematically delineated the vital role of macrophages in mediating the CA promoting thrombus resolution.

## Materials and methods

### Materials

The main materials used in this study were caffeic acid (Molecular formula: C_9_H_8_O_4_, Molecular weight: 180.16, purity: 98%, Shanghai Tauto Biotech CO., LTD, Shanghai, China); clodronate liposomes and PBS liposomes (Liposoma, Netherlands); RIPA lysis buffer and gelatin zymography analysis Kit (Real-Times Biotechnology Co.,Ltd. Beijing, China); Granulocyte-macrophage colony-stimulating factor (GM-CSF) (Peprotech, USA); RPMI 1640 medium (Gibco, USA), LPS (Sigma-Aldrich, USA), IL-4 (Peprotech, USA), IL-13 and IFN-γ (MedChemExpress, China). Other reagents were of analytical grade.

### Animals

C57BL/6J male mice weighing 18 ~ 22g were obtained from SLRC Laboratory Animal Lid (Shanghai, China). Wild-type mice and Nrf2 gene-deficient mice weighing 18 ~ 22g (kindly donated by Professor Li-li Ji of Shanghai University of Traditional Chinese Medicine). All mice were kept under an automated 12-hour dark-light cycle at a controlled temperature of 22°C ± 2°C and a relative humidity of 50%~60% with free access to standard dry diet and tap water *ad libitum*.

All animals received humane care and experimental procedures were performed in accordance with the guidelines of the Shanghai University of Traditional Chines Medicine for health and care of experimental animals. The protocol was reviewed and approved by the Experimental Animal Ethical Committee of Shanghai University of Traditional Chinese Medicine (Permit Number: PZSHUTCM2408210001).

### Stasis-induced model of deep vein thrombosis

DVT was induced by inferior vena cava (IVC) ligation as described previously.^25^ In brief, mice were anesthetized using isoflurane (oxygen delivered at 0.5 L/min with 3% isoflurane for induction and 1.5% isoflurane for maintenance) for 15 min. After a midline abdominal incision was made, inferior vena cava was exposed carefully and the lumbar side branches of the IVC were cauterized. Then the vena cava was ligated with 6-0 silk suture immediately below the the level of the renal veins. Then the muscle layer and epidermal layer were sutured. Finally, sterile isotonic saline was administered for volume substitution. Sham-operated mice were undergoing laparotomy with only exposition of IVC without ligation. After recovery from anesthesia, mice were given free access to food and water. Mortality was assessed daily.

Caffeic acid was dissolved in saturated sodium bicarbonate. Caffeic acid was prepared and used on daily basis. In the caffeic acid experiment, mice were treated with caffeic acid (20, 40, 80 mg/kg/day) by gavage 3 days prior to the IVC ligation and continued daily to either day 3 or day 7 or day 14 after surgery.

To investigate the role of macrophages in the thrombus resolution treated with caffeic acid, DVT mice were divided randomly into three groups: CA + PBS liposomes group, CA + chlorophosphate liposomes group, and CA + chlorophosphate liposomes + BMDM group. Bone marrow-derived macrophages (BMDMs) from mice were intraveously infused into macrophage-depleted mice (1×10^6^ cells per mouse) on the day 3 post IVC ligation.

To investigate the role of Nrf2 in the thrombus resolution treated with caffeic acid, DVT mice were divided randomly into two groups: CA + WT group and CA + *Nrf2^-/-^* group.

The mice were then sacrificed at day 3, 7, or 14 and the thrombosed vena cava distal to the renal veins up to the iliac bifurcation was excised, length measured, and weighed in milligrams using an analytical balance. At harvest, thrombus and vascular wall tissues were collected for histological and subsequent molecular analyses.

### Depletion of macrophages in vivo

Mouse macrophages were depleted as described previously.^26^ Briefly, 10 mL/kg of either clodronate liposomes or control liposomes were administered to the mouse through tail vein injection once at day 2 after IVC ligation. However, since myeloid cells are replenished quickly, evaluation of longer time-points required continuous injection with clodronate liposomes every three days until the end of the experiment. The thrombi were harvested at day 7 after surgery.

### Bone marrow-derived macrophages isolation and culture

Bone marrow-derived macrophages (BMDMs) were isolated from the femurs and tibiae of mice as previously described.^27^ Erythrocytes were removed through a 70 μm cell strainer and subjected to red blood cell lysing buffer and resuspended in RPMI 1640 medium. Cells were cultured and treated with macrophage colony-stimulating factor (GM-CSF, 50 ng/mL) for 6 days. Half volume medium change on day 3. On the seventh day, nonadherent cells were removed, and the medium was replaced. BMDM cells were pretreated with caffeic acid (50 µmol/L) for 12 h, and then exposed to LPS (100 ng/mL) and IFN-γ (50 ng/mL) for 24 h to induce classical macrophage differentiation (M1 polarization), or to recombinant IL-4 (10 ng/mL) and IL-13 (10 ng/mL) for 24 hours to induce alternatively activated macrophage differentiation (M2 polarization). After incubation, BMDMs were applied to subsequent analyses

### Histopathological Evalution

Segments of thrombus and vascular wall tissues were fixed in 10% phosphate-buffered saline buffered formalin and embedded in paraffin. Approximately 4 serial sections (4 μm thick) were cut, placed on glass slides. One slide was stained with hematoxylin and eosin (H&E). The other two slides were prepared for Masson’s trichrome staining, immunofluorescence. All images were acquired using the OLYMPUS IX71 microscope. Collagen fibers were dyed blue. The percentage of the collagen content in the thrombus was quantified through computerized analysis using Image-Pro Plus version 5.0.

### Immunofluorescence assay

Thrombus and vascular wall tissues were fixed and embedded in paraffin and then sectioned in 4 μm slices. Subsequently, the samples were incubated with primary antibody for CD68, iNOS, Arg-1, CD31 in 1% BSA/PBS overnight at 4℃, and then washed and incubated with secondary antibody or fluorescent secondary antibody for 1 hour at room temperature. The samples were washed, dried, mounted in medium containing DAPI for nuclear DNA staining, and imaged on an Olympus IX71 microscope.

### Immunoblotting

Total proteins were isolated from thrombus and vascular wall tissue or BMDM cells homogenates. Nuclear proteins were isolated by using the ExKine™ nuclear protein extraction Kit (Abbkine Scientific Co.,Ltd, Wuhan, China). After sonication and centrifugation, the supernatant was used for determination of protein concentrations by the Bradford method. Equal amounts of protein were resolved in SDS-PAGE and transferred to nitrocellulose membrane. The proteins of interest were detected using specific antibodies against CD68 (Affinity, 1:1000), CD31 (Cell Signaling, 1:1000), iNOS (Affinity, 1:1000), Arginase-1 (Cell Signaling, 1:1000), IL-1β (ABclonal, 1:1000), TNF-α (Cell Signaling, 1:1000), BAX (ABclonal, 1:1000), Bcl-2 (Proteintech, 1:5000), cleaved Caspase-1 (Affinity, 1:1000), TGF-β (Affinity, 1:1000), VEGFA (ABclonal, 1:1000), Nrf2 (ABclonal, 1:1000), Keap-1 (Servicebio, 1:500), Lamin B1 (ABclonal, 1:5000) and GAPDH (Proteintech, 1:5000). Bands were visualized and quantified with Odyssey Infrared Imaging System (LI-COR, Lincoln, NE, USA). All immunoblotting experiments were repeated at least three times.

### Gelatin Zymography assay

Thrombus protein lysates were prepared using EDTA-free RIPA lysis buffer. Gelatin zymography analysis Kit was used. Equal concentrations of protein were loaded onto zymogram gels and run at 125 V constant for 90 minutes. The gels were then developed and incubated in 1×Zymogram Renaturing Buffer and 1× Zymogram Developing Buffer per the manufacturer’s instructions. Gels were then stained with FastBlue protein stain and imaged. Intensities of the bands were quantified and calculated using Image J software.

### RNA-sequencing analysis

BMDM were isolated, cultured and induced by IFN-γ+LPS as described above. Total RNA was extracted from caffeic acid stimulated and unstimulated cells, frozen, and sent for sequencing (Majorbio Technologies Inc., Shanghai, China). RNA quality and integrity were assessed using the Agilent 2100 Bioanalyzer (Agilent Technologies, USA). Samples with an RNA integrity number (RIN) > 7.0 were selected for further analysis. Library preparation was performed using the Illumina TruSeq Stranded mRNA Library Prep Kit, following the manufacturer’s protocol. The prepared libraries were sequenced on an Illumina platform in paired-end mode with a read length of 150 bp. Raw sequencing reads were processed using FastQC for quality control, and adapter sequences were trimmed using Trimmomatic. Clean reads were aligned to the mouse reference genome (GRCm39) using HISAT2, and gene expression levels were quantified using featureCounts. Differential gene expression analysis was performed using DESeq2, with an adjusted p-value < 0.05 considered statistically significant.

### Functional enrichment analysis

For functional enrichment analysis, gene set enrichment analysis (GSEA) was conducted using the GSEA software (Broad Institute), with the KEGG and Gene Ontology (GO) gene sets used as references. The enrichment results were considered significant when FDR < 0.25.

### Statistical Analysis

All the data were expressed as mean ± SEM. Unpaired Student’s t-test was used for comparison of data from two groups. One way analysis of variance (ANOVA) was used for comparison of data involving more than two groups. *P*<0.05 was considered statistically significant. The diagrams and statistical analyses were performed in GraphPad Prism 8.

## Results

### Caffeic acid promoted the resolution of DVT

It was reported that caffeic acid significantly prolonged thrombosis latency in murine cerebral venules subjected to chemical stimulation and reduced the thrombus-to-vessel area ratio.^23^ To evaluate its potential therapeutic effects on thrombus resolution, caffeic acid was pre-administering three days before inferior vena cava (IVC) ligation. The protocol is shown in **Figure 1A** and schematic diagram of stasis-induced mice deep vein thrombosis model in **Figure 1B**. Thrombus burden was assessed at 3, 7, and 14 days after IVC ligation. The representative diagrams of the thrombus tissue samples were shown in **Figure 1C**. Macroscopically, IVC thrombus showed progressive changes in size and composition from day 3 to day 14. The results of thrombus length and thrombus weight showed a gradual decrease in thrombus burden, with a significant decrease from day 3 to day 14. It was known that the period from day 1 to day 3 after modeling or thrombosis was considered the thrombus formation period, and the early stage of thrombus resolution was considered from day 4 to day 7 after thrombosis and the middle and late stages of thrombus resolution were considered from day 8 after thrombosis.^7,8^ Amazingly, as shown in **Figure 1D, 1G**, thrombus length and thrombus weight did not change in caffeic acid group compared with DVT group at 3 days, suggesting that caffeic acid did not affect venous thrombus formation. However, thrombus length and thrombus weight were reduced in caffeic acid (40 mg/kg) group before IVC ligation at 7 and 14 days compared with DVT group treated with vehicle (**Figure 1E, 1F, 1H and 1I**). Hence, caffeic acid (40 mg/kg) for continuous 3 days before IVC ligation was used in subsequent experiments. These results suggested that caffeic acid promoted venous thrombus resolution in the early stage.

**Figure 1.**
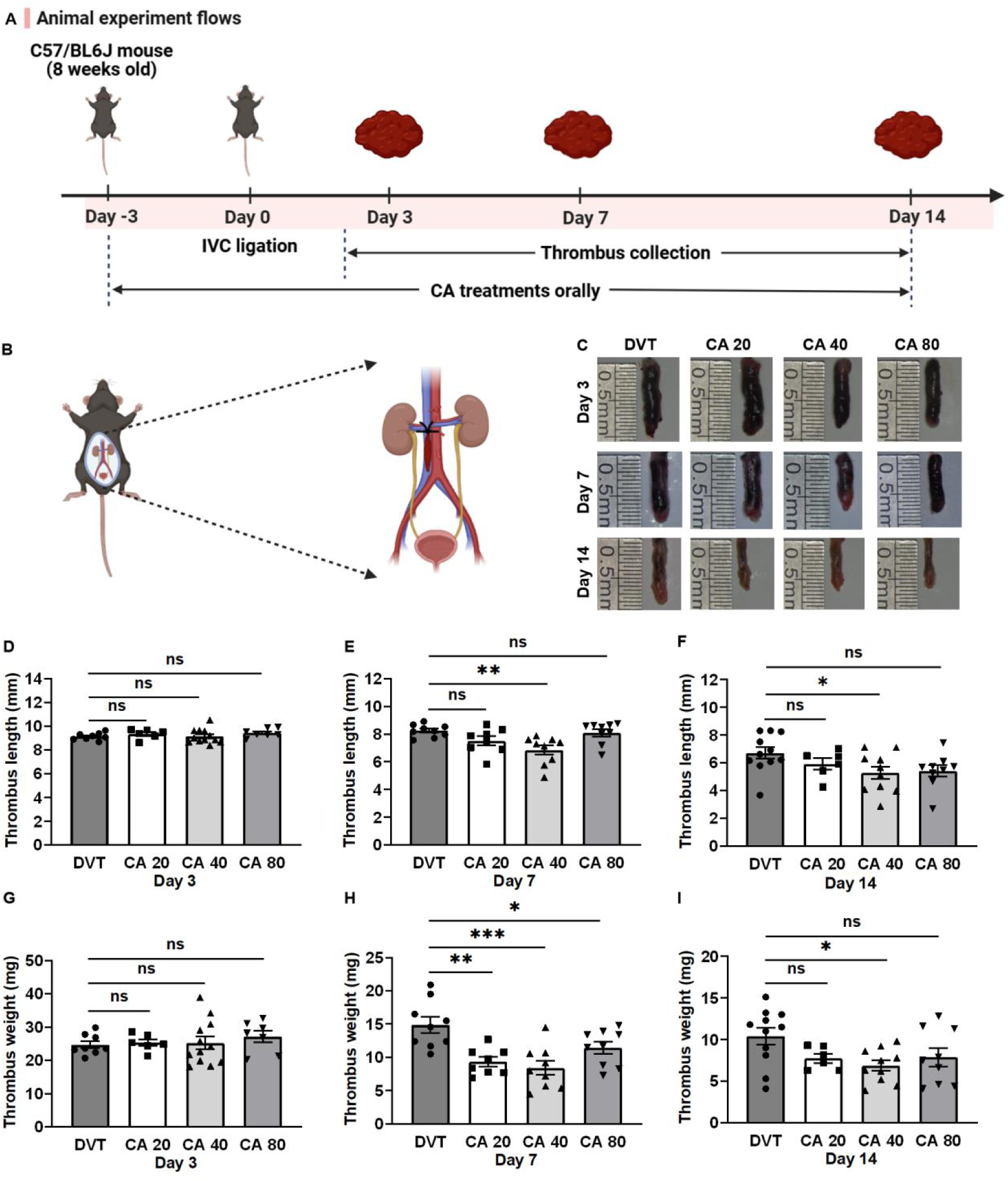
Caffeic acid enhanced the resolution of DVT. (A) The flow chart. (B) Schematic diagram of IVC ligation. (C) Macroscopic picture of thrombus in mice at 3, 7, and 14 days post IVC ligation. (D-I) Thrombus lengths and weight from mice at day 3, 7 and 14 after IVC ligation treated with caffeic acid (40 mg/kg). *n*=6-12 per group. All values represented the mean ± SEM. \**P*<0.05, \*\**P*<0.01, \*\*\**P*<0.001, *ns,* no significance; CA, caffeic acid; DVT, deep vein thrombosis.

### Caffeic acid promoted collagenolysis in thrombus

As fibrosis and collagen remodeling occur in the inflammatory vascular remodeling of venous thrombi during venous thrombus resolution, the collagen content in thrombus at the day 7 and day 14 was evaluated by histopathological analysis. Hematoxylin-Eosin staining showed that thrombus area was not change at day 3, 7, and 14 in caffeic acid group compared with DVT group (**Figure 2A**). However, Masson’s Trichrome Stain showed that intrathrombotic collagen was significantly reduced at the day 14 in caffeic acid group compared with DVT group (**Figure 2A**). It was reported that the matrix metalloproteinases 2 and 9 (MMP-2 and MMP-9) play key roles in collagenolysis and matrix remodeling during thrombus resolution.^28,29^ As shown in **Figure 2B, 2C,** the MMP-9 activity was not different between caffeic acid group and DVT group, while the MMP-2 activity was found to be significantly elevated in the thrombus of caffeic acid group compared with DVT group at day 14 after IVC ligation.

**Figure 2.**
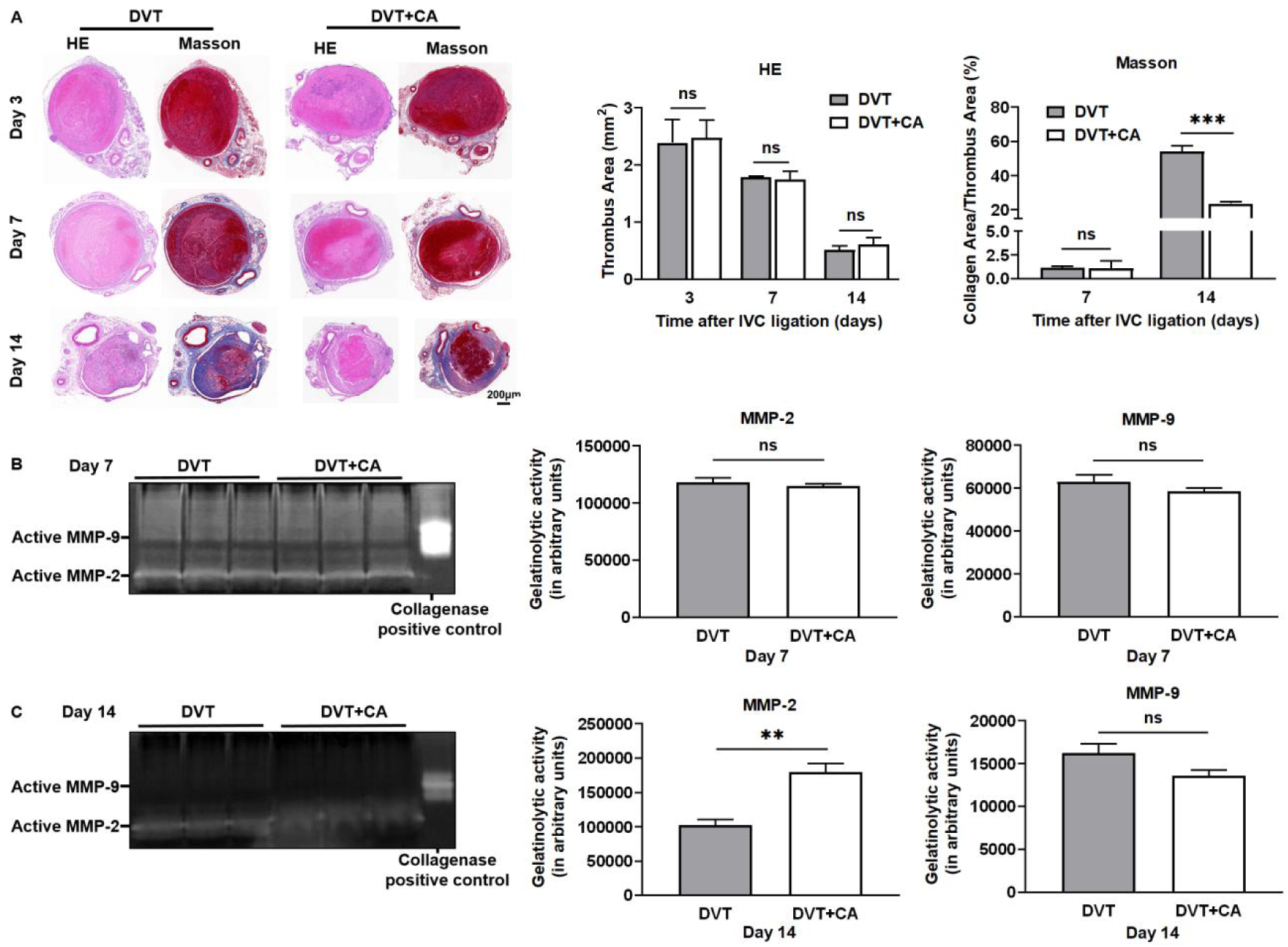
Caffeic acid promoted collagenolysis in thrombus. (A) Representative images of thrombus sections stained with HE and Masson and quantitative comparison of thrombus area and collagen area/thrombus area from mice at day 3, 7 and 14 after IVC ligation treated with caffeic acid (40 mg/kg). Scale bars are 200 μm. Data are presented as the mean ± SEM; *\*\*\*P<0.001*, *n*=3 per group. Representative images of active MMP-2 and active MMP-9 in thrombus of mice at day 7 (B) and day 14 (C) after IVC ligation treated with caffeic acid (40 mg/kg) and representative gel images of intrathrombotic MMP-2 and MMP-9 activities in venous thrombus samples from DVT measured by gelatin gel zymography. *n*=3, respectively. The experiment was repeated at least three times. Gel images were subjected to semiquantitative analysis as described in “Methods.” All values represented the mean ± SEM. *\*\*P<0.01*. *ns*, no significance; HE, hematoxylin and eosin; CA, caffeic acid; MMP-9, matrix metalloproteinase-9; MMP-2, matrix metalloproteinase-2; DVT, deep vein thrombosis.

### Caffeic acid promoted macrophage infiltration into thrombus

It was demonstrated that the increase of macrophage numbers or monocyte recruitment into the thrombus could improve VT resolution and recanalization.^30^ Firstly, the effect of caffeic acid on macrophage infiltration and macrophage polarization in thrombus was observed by immunoblotting. As shown in **Figure 3A**, caffeic acid significantly increased the expression of the macrophage marker CD68 and inhibited the expression of the macrophage M1 polarization marker iNOS, but did not affect the expression of the macrophage M2 polarization marker Arg-1 in thrombus. Similarly, the immunofluorescence double staining showed that caffeic acid significantly increased the expression of the macrophage marker CD68 and inhibited the expression of the macrophage M1 polarization marker iNOS, but did not affect the expression of the macrophage M2 polarization marker Arg-1 in thrombus. (**Figure 3B-C**). Furthermore, we also observed on the immunofluorescence double staining that the macrophages in the DVT group were mainly distributed around the periphery of venous thrombus, while the macrophages in the caffeic acid group were also expressed in the center of venous thrombus. It indicated that caffeic acid can increase the content and infiltration of macrophages in venous thrombus.

**Figure 3.**
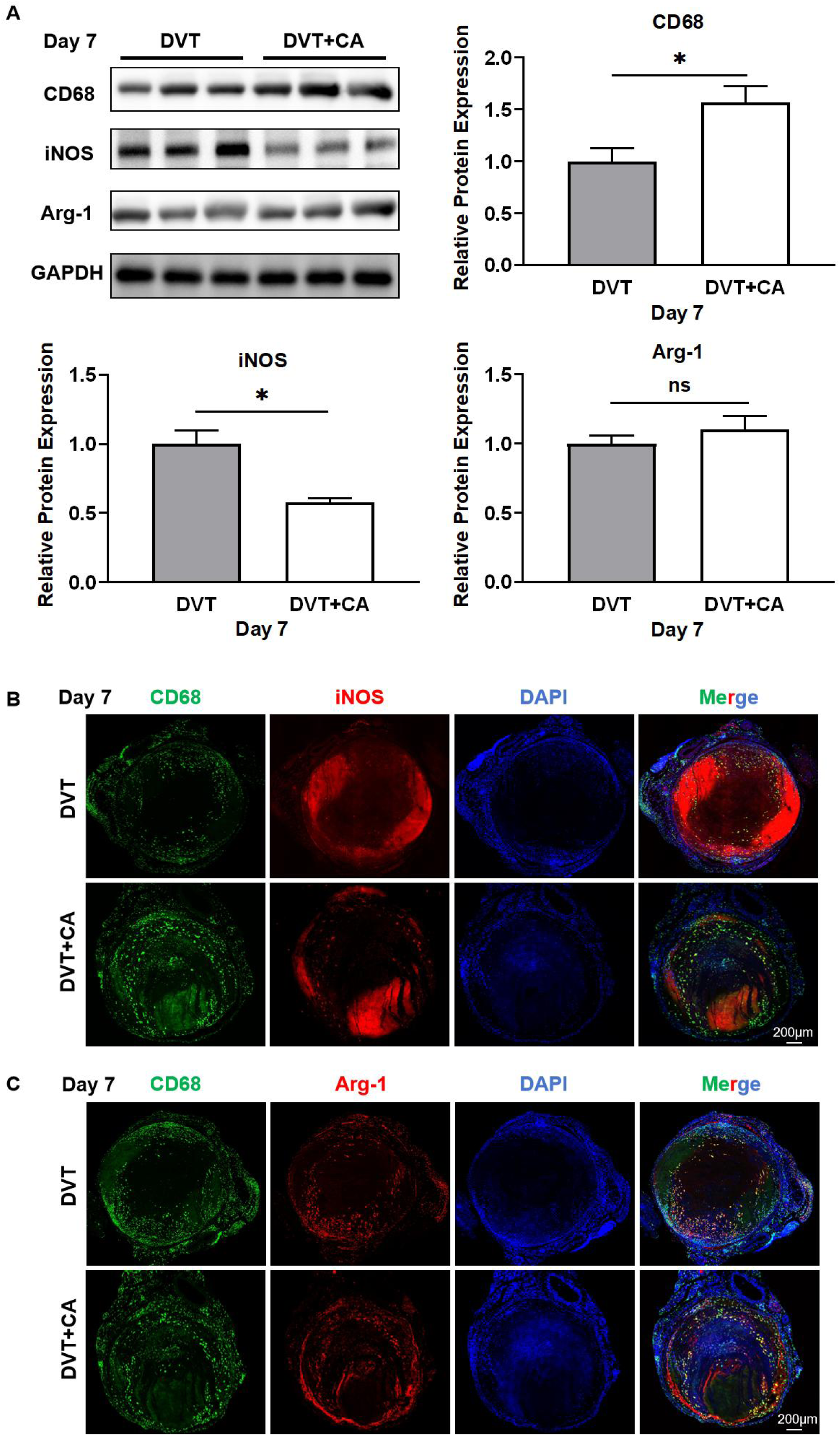
Caffeic acid promoted macrophage infiltration in thrombus. (A) Representative Western blot for CD68, iNOS and Arg-1 expression in thrombus tissue from mice at day 3, 7 and 14 after IVC ligation treated with caffeic acid (40 mg/kg) and densitometric analysis (left). The experiment was repeated at least three times. GAPDH served as the loading control. Data are presented as the mean±SEM. \**P*<0.01. (B) Representative images of immunofluorescent staining for CD68 (green) and iNOS (red) in thrombus tissues. (C) Representative images of immunofluorescent staining for CD68 (green) and Arg-1 (red) in thrombus tissues. The localization of the nucleus was detected by DAPI staining (blue). *n*=3 per group. Scale bars are 200 μm, CD68, cluster of differentiation 68; iNOS, inducible nitric oxide synthase; Arg-1, Arginine-1; DAPI, 4’,6-diamidino-2-phenylindole; *ns,* no significance; CA, caffeic acid; DVT, deep vein thrombosis.

The results indicated that caffeic acid increased macrophage content and inhibited macrophage M1 polarization, but did not affect macrophage M2 polarization in thrombus.

### Caffeic acid inhibited inflammation in thrombus

The process of venous thrombus resolution is associated with a series of alterations in the expression of inflammatory cytokines.^12^ Therefore, firstly, inflammatory factor interleukin-1β (IL-1β) in thrombotic tissues was detected by immunoblotting. As shown in **Figure 4A**, caffeic acid significantly inhibits the expression of IL-1β in thrombus. However, caffeic acid did not affect the expression of apoptotic markers BAX and Bcl-2, and also did not affect the expression of Casepase-1 and TGF-β.

**Figure 4.**
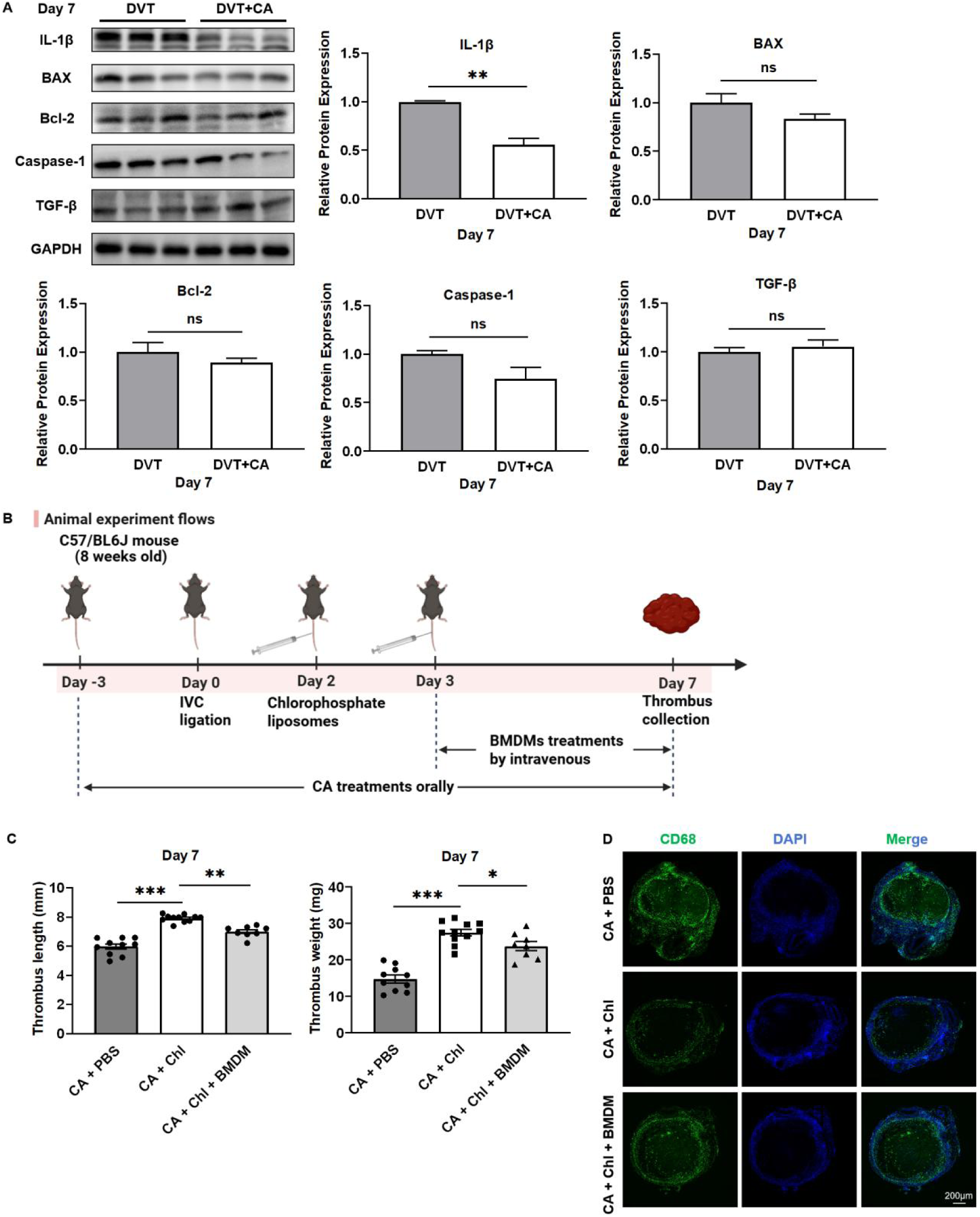
Caffeic acid inhibited inflammation in thrombus. (A) Representative Western blot for IL-1β, BAX, Bcl-2, Caspase-1 and TGF-β expression in thrombus tissue from mice at day 3, 7 and 14 after IVC ligation treated with caffeic acid (40 mg/kg) and densitometric analysis (right). The experiment was repeated at least three times. GAPDH served as the loading control. Data are presented as the mean±SEM. \*\**P*<0.01. (B) The flow chart. Thrombus lengths (C) and weight (D) in chlorophosphate liposomes-treated mice at day 7 after IVC ligation. *n*=8-11 per group. All values represented the mean ± SEM. \**P*<0.05, \*\**P*<0.01, \*\*\**P*<0.001, (E) Representative images of immunofluorescent staining for CD68 (green) in thrombus tissues. The localization of the nucleus was detected by DAPI staining (blue). *n*=3-5 per group. Scale bars are 200 μm, BMDM, bone marrow-derived macrophages; DAPI, 4’,6-diamidino-2-phenylindole; *ns,* no significance; CA, caffeic acid; DVT, deep vein thrombosis.

Macrophages are the main inflammatory cells in thrombus.^31^ To demonstrate that caffeic acid inhibited inflammation to promote thrombus resolution via macrophages, the chlorophosphate liposome was used to depletion of macrophages. The experiment was divided into three groups (CA + PBS liposomes, CA + Chlorophosphate liposomes, and CA + Chlorophosphate liposomes + BMDM) and the the protocol was shown in **Figure 4B**. As shown in **Figure 4C-D**, thrombus length and thrombus weight were significantly increased after macrophage deletion by chlorophosphate liposomes (CA + chlorophosphate liposomes group) compared with the CA + PBS liposomes group. However, intravenous injection of BMDM treatment (CA + clophosphate liposomes+BMDM group) decreased thrombus length and thrombus weight compared with the CA + chlorophosphate liposomes group. Furthermore, to verify that macrophages were deleted by clophosphate liposomes, immunofluorescence double staining method was used. As shown in **Figure 4E**, little macrophages were observed in the CA + clophosphate liposomes group compared to the CA + PBS liposomes group, whereas the number of macrophages was restored in the CA + clophosphate liposomes + BMDM treatment group.

### Caffeic acid promoted neovascularizationin thrombus

It was found that promoting neovasculation is one of main way through which macrophages promote thrombus resolution.^32^ Therefore, vascular endothelial growth factor (VEGF) and CD31, the angiogenic markers, were detected by immunoblotting. It was showed that caffeic acid significantly increased the expression of VEGFA and CD31 in thrombus tissue **(Figure 5A)**. Similarly, immunofluorescence double staining results showed that caffeic acid increased the expression of CD31 in thrombus **(Figure 5B**). The results suggest that caffeic acid promoted neovascularization in the thrombus, thereby facilitating thrombus resolution. It is worth noting that we observed that neovascularization was mainly near macrophages, suggesting that it might be the VEGFA secreted by macrophages that promoted angiogenesis, thereby facilitating the thrombus resolution.

**Figure 5.**
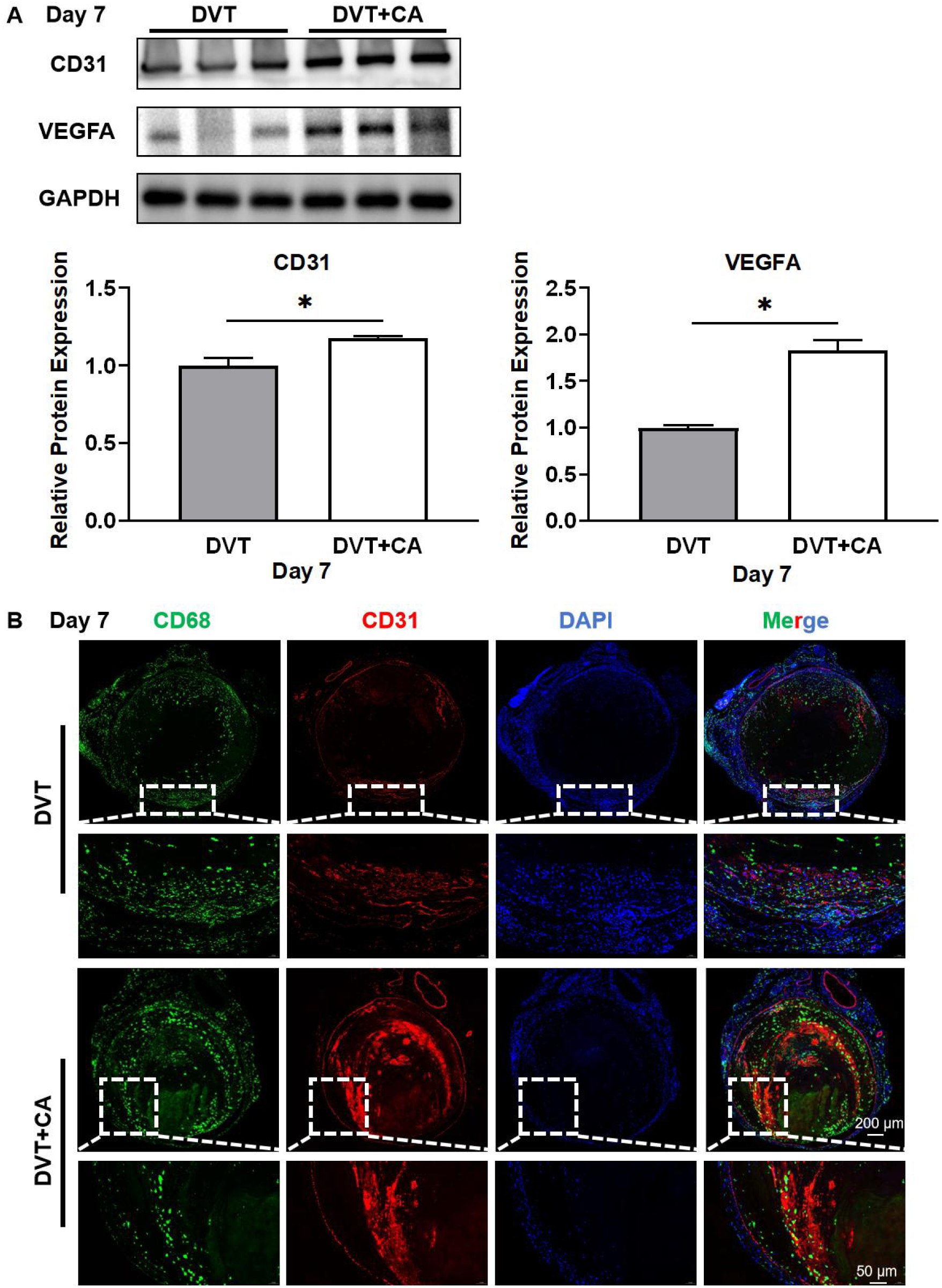
Caffeic acid promoted neovasculation in thrombus. (A) Representative Western blot for CD31 and VEGFA expression in thrombus tissue from mice at day 3, 7 and 14 after IVC ligation treated with caffeic acid (40 mg/kg) and densitometric analysis. The experiment was repeated at least three times. GAPDH served as the loading control. Data are presented as the mean±SEM. \**P*<0.05. (B) Representative images of immunofluorescent staining for CD68 (green) and CD31 (red) in thrombus tissues. The localization of the nucleus was detected by DAPI staining (blue). *n*=3 per group. Scale bars are 200 μm, CD31, Platelet endothelial cell adhesion molecule-1; VEGFA, vascular endothelial growth factor A;

### Caffeic acid inhibits macrophage M1 polarization in BMDMs induced by LPS and IFN-γ

In order to determine the effect of caffeic acid on macrophage polarization, BMDM cells were exposed to caffeic acid (50 µM) for 12 h and subsequently cultured with LPS (100 ng/mL) and IFN-γ (50 ng/mL) or IL-4 (10 ng/mL) and IL-13 (10 ng/mL) for 24 h. The protein expression of the iNOS and IL-1β was quantified by immunoblotting and qRT-PCR. The protocol is shown in **Figure 6A**. As shown in **Figure 6B**, caffeic acid inhibited the expression of iNOS, IL-1β and casepase-1 in BMDMs induced by LPS + IFN-γ, suggesting that caffeic acid inhibited M1 polarization in macrophages. However, the expression of BAX, Bcl-2, TGF-β, CD31 and VEGFA in BMDMs induced by LPS + IFN-γ did not alter after treated with caffeic acid. As shown in **Figure 6C**, caffeic acid did not influence the expression of Arg-1 in BMDMs induced by IL-4 and IL-13, suggesting that caffeic acid did not promote M2 polarization in macrophages.

**Figure 6.**
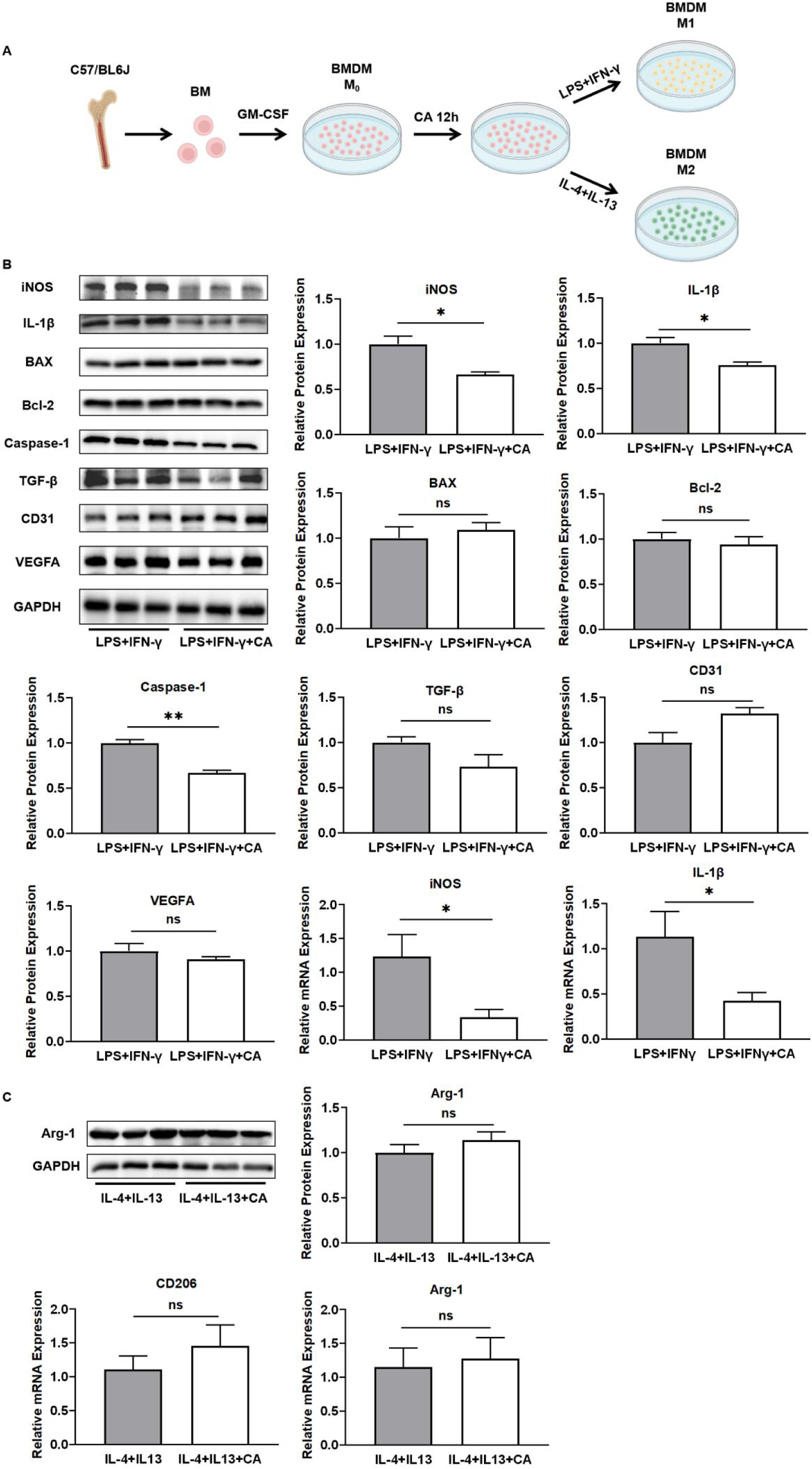
Caffeic acid inhibited macrophage M1 polarization in BMDMs induced by LPS and IFN-γ *in vitro*. (A) The flow chart. (B) Representative Western blot for iNOS, IL-1β, BAX, Bcl-2, Caspase-1, TGF-β, CD31 and VEGFA expression and representative RT-qPCR for iNOS, IL-1β expression in bone marrow-derived macrophages induced by LPS and IFN-γ for 24 hours *in vitro* and densitometric analysis. The experiment was repeated at least three times. GAPDH served as the loading control. Data are presented as the mean±SEM. \**P*<0.05, \*\**P*<0.01. (C) Representative Western blot for Arg-1 expression and representative RT-qPCR for CD206, Arg-1 expression in bone marrow-derived macrophages induced by IL-4 and IL-13 *in vitro* and densitometric analysis. The experiment was repeated at least three times. GAPDH served as the loading control. Data are presented as the mean±SEM. LPS, lipopolysaccharide; IFN-γ, interferon-γ; IL-1β, interleukin-1β; IL-4, interleukin-4; IL-13, interleukin-13; TGF-β, transforming growth factor-β; Arg-1, Arginine-1; *ns,* no significance; CA, caffeic acid.

### Caffeic acid attenuated inflammatory activity through Nrf2 pathways both in BMDMs induced by LPS and IFN-γ and in thrombus tissue from mice after IVC ligation

To investigate the potential mechanism of action of caffeic acid in promoting thrombus resolution, BMDMs pretreated by caffeic acid (50 µM) were induced by LPS and IFN-γ, followed by RNA sequencing of BMDMs. The results showed that the anti-oxidative stress-related genes Hmox1, Nqo1, Gclm, Gss, Gclc, Txnrd1, Prdx1, and Slc7a11 were significantly up-regulated in BMDM after 12 h of caffeic acid intervention (**Figure 7A**), which were mainly regulated by nuclear factor erythroid 2-related factor 2 (Nrf2) signaling pathway, which is closely related to cellular defense against oxidative stress. In addition, heat map analysis (**Figure 7B**) showed that the expression levels of antioxidant genes were significantly increased after CA intervention, while inflammation-related genes (e.g., Il6, Ccl7, Ccl12, and Cxcl10) were significantly down-regulated, suggesting that CA may inhibit inflammatory responses through the Nrf2 pathway. To further confirm the effect of CA on the Nrf2 signaling pathway, we performed functional enrichment of differential genes using Gene Set Enrichment Analysis (GSEA). The analysis results showed that Nrf2 related pathways (e.g. Glutathione metabolism, Nrf2-ARE regulation, Oxidative stress response) were significantly activated with high enrichment scores (NES > 2, *P* < 0.002) after CA treatment (**Figure 7C**). The activation of these pathways further supports that CA inhibits M1-type polarization and promotes M2-type polarization through the enhancement of Nrf2-mediated antioxidant stress response, thereby reducing the inflammatory response in BMDM.

**Figure 7.**
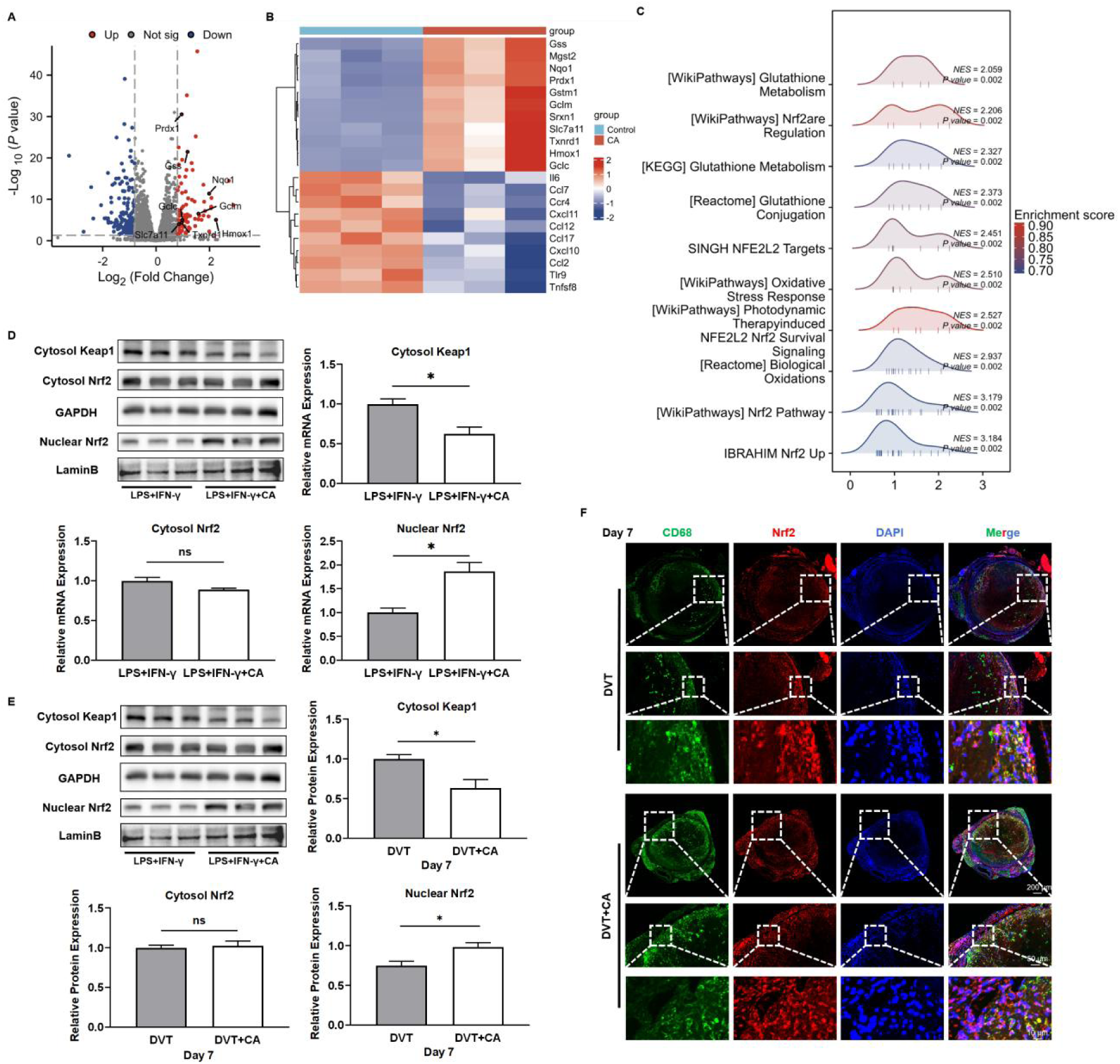
Caffeic acid attenuated inflammatory activity through Nrf2 pathways both in BMDMs induced by LPS and IFN-γ and in thrombus tissue from mice after IVC ligation. Caffeic acid regulated gene expression in BMDM through Nrf2 signaling pathway. (A) Volcano plot (Volcano plot) showing the up-regulated (red) and down-regulated (blue) differential genes in BMDM after CA intervention, in which antioxidant-related genes Hmox1, Nqo1, Gclm, Gss, Gclc, Txnrd1, Prdx1, Slc7a11 were significantly up-regulated. (B) Heatmap demonstrating the gene expression of BMDM in CA-treated and control groups. (C) Results of GSEA enrichment analysis, NRF2-related pathways (e.g. Glutathione metabolism, Nrf2 are Regulation) were significantly up-regulated after CA intervention. *n* = 3 per group. (D) Representative Western blot for nuclear Nrf2, Lamin B, cytosol Nrf2 and cytosol Keap1 expression in bone marrow-derived macrophages induced by LPS and IFN-γ for 24 hours *in vitro* and densitometric analysis. The experiment was repeated at least three times. GAPDH served as the loading control. Data are presented as the mean±SEM. \**P*<0.05. (E) Representative Western blot for nuclear Nrf2, Lamin B, cytosol Nrf2 and cytosol Keap1 expression in thrombus tissues from mice at day 7 after IVC ligation treated with caffeic acid (40 mg/kg) and densitometric analysis. The experiment was repeated at least three times. GAPDH served as the loading control. Data are presented as the mean±SEM. \**P*<0.05. (F) Representative images of immunofluorescent staining for CD68 (green) and Nrf2 (red) in thrombus tissues. The localization of the nucleus was detected by DAPI staining (blue). *n*=3 per group. Scale bars are 200, 50 and 10 μm. Nrf2, nuclear factor erythroid 2-related factor 2;

It was found that Nrf2 is involved in regulating oxidative stress, inflammation, autophagy, and immune response. Normally, Nrf2 is located in the cytoplasm and bound to the E3 ligase adaptor protein, Kelch-like ECH-associated protein 1 (Keap1),^33^ but Nrf2 dissociates from Keap1 and enters the nucleus and activates antioxidant genes during oxidative stress.^34,35^ Hence, whether caffeic acid promotes the translocation of Nrf2 to the nucleus was verified through *in vitro* and *in vivo* experiment. As shown in **Figure 7D**, BMDM cells were exposed to caffeic acid (50 µM) for 12 h, and then incubated with LPS (100 ng/mL) and IFN-γ (50 ng/mL) for 24 h. It was showed that caffeic acid inhibited the expression of Keap-1 and promoted the translocation of Nrf2 into the nucleus. Similarly, as shown in **Figure 7E**, caffeic acid inhibited the expression of Keap1 and promoted the translocation of Nrf2 into the nucleus in thrombotic tissue of mice compared with the DVT group. Similarly, the immunofluorescence double staining showed that caffeic acid significantly promoted the translocation of Nrf2 into the nucleus in thrombotic tissue of mice compared with the DVT group **Figure 7F**.

### Nrf2 deletion abgrogates the pro-resolving effects of caffeic acid

To confirm the key role of Nrf2 in promoting deep vein thrombus resolution through caffeic acid, we used Nrf2 gene-deficient mice. The experiment was divided into two groups: CA + WT and CA + *Nrf2^-/-^*. As shown in **Figure 8A**, compared with the CA + WT group, the length and weight of venous thrombus significantly increased after Nrf2 gene deletion (CA + *Nrf2^-/-^* group). These results indicated that Nrf2 plays a key role in promoting deep vein thrombus resolution through caffeic acid.

**Figure 8.**
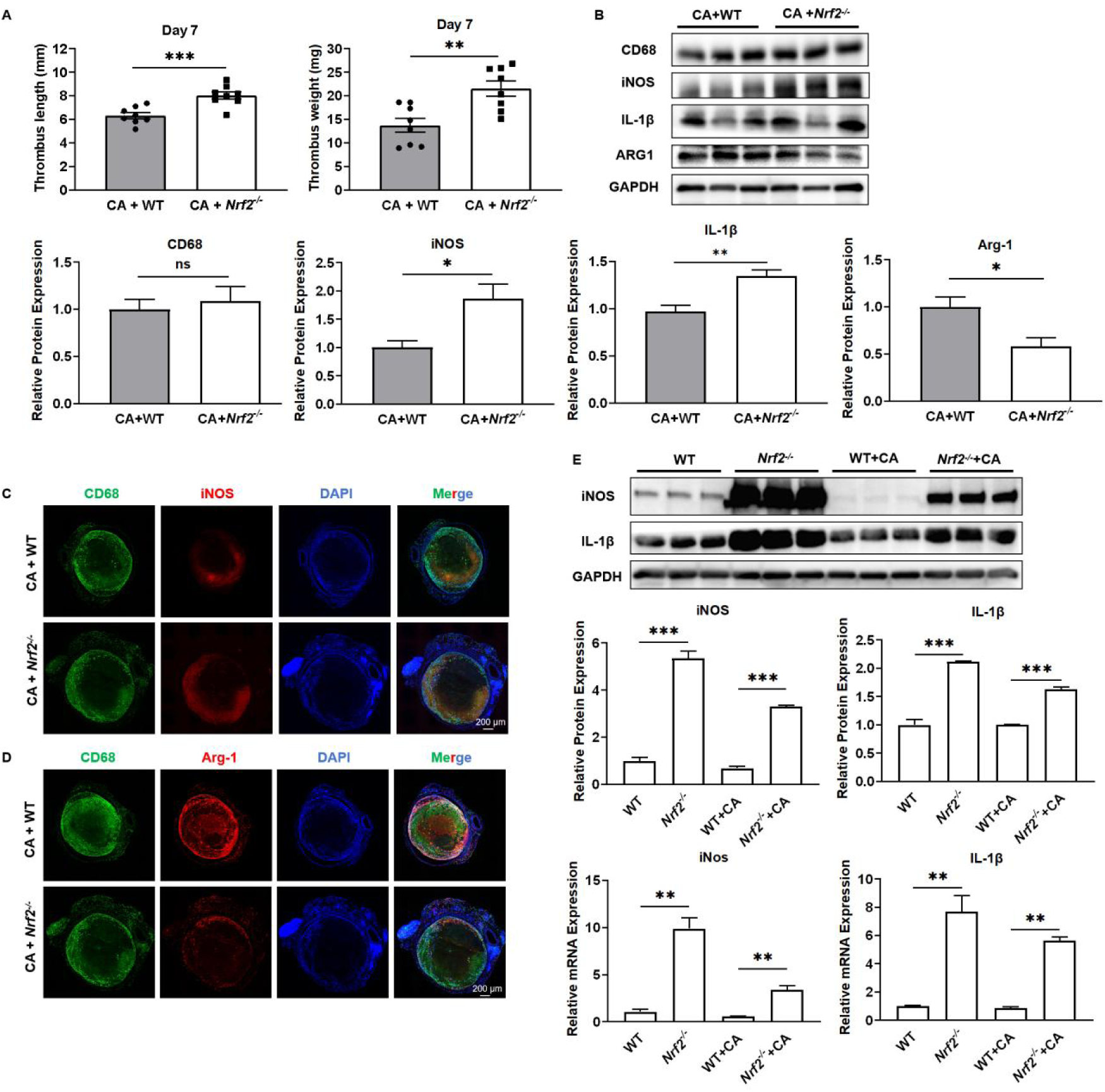
Nrf2 deletion abrogates the pro-resolving effects of caffeic acid. (A) Thrombus lengths and weight from *Nrf2^-/-^* mice at day 7 after IVC ligation treated with caffeic acid (40 mg/kg). n=8 per group. All values represented the mean ± SEM. *\*P<0.05*, *\*\*P<0.01*, *\*\*\*P<0.001*, *ns*, no significance; CA, caffeic acid; DVT, deep vein thrombosis. (B) Representative Western blot for CD68, iNOS, IL-1β and Arg-1 expression in thrombus tissue from *Nrf2^-/-^* mice at day 7 after IVC ligation treated with caffeic acid (40 mg/kg) and densitometric analysis (right). The experiment was repeated at least three times. GAPDH served as the loading control. Data are presented as the mean ± SEM. *\*P<0.05*, *\*\*P<0.01*. (C) Representative images of immunofluorescent staining for CD68 (green) and iNOS (red) in *Nrf2^-/-^* mice thrombus tissues. (D) Representative images of immunofluorescent staining for CD68 (green) and Arg-1 (red) in *Nrf2^-/-^* mice thrombus tissues. The localization of the nucleus was detected by DAPI staining (blue). *n*=3 per group. Scale bars are 200 μm, CD68, cluster of differentiation 68; iNOS, inducible nitric oxide synthase; Arg-1, Arginine-1; DAPI, 4’,6-diamidino-2-phenylindole; ns, no significance; CA, caffeic acid; DVT, deep vein thrombosis. (E) Representative Western blot for iNOS and IL-1β expression and representative qRT-PCR for iNOS and IL-1β expression in bone marrow-derived macrophages induced by LPS and IFN-γ for 24 hours *in vitro* and densitometric analysis. The experiment was repeated at least three times. GAPDH served as the loading control. Data are presented as the mean±SEM. \**P*<0.05, \*\**P*<0.01.

The effects of Nrf2 gene deletion on macrophage infiltration and macrophage polarization in caffeic acid promoted venous thrombolysis were observed by Western blotting. As shown in **Figure 8B**, compared with the CA + WT group, Nrf2 gene deletion (CA + *Nrf2^-/-^* group) did not affect the expression of macrophage marker CD68 in venous thrombus, but promoted the expression of macrophage M1 polarization marker iNOS and inflammatory factor IL-1β, and inhibited the expression of the M2 polarization marker Arg-1 in macrophages. Similarly, immunofluorescence double staining showed that, compared with the CA + WT group, Nrf2 gene deletion (CA + *Nrf2^-/-^* group) did not affect the expression and infiltration of macrophage marker CD68 in venous thrombus, but promoted the expression of macrophage M1 polarization marker iNOS, and inhibited the expression of the M2 polarization marker Arg-1 in macrophages in **Figures 8C and 8D**. Furthermore, we also founded that the deletion of the Nrf2 gene promoted the expression of the M1 polarization marker iNOS and the inflammatory factor IL-1β in BMDM cells stimulated by LPS (100 ng/mL) and IFN-γ (50 ng/mL) for 24 h in **Figures 8E**.

## Discussion

In the current study, we demonstrated that caffeic acid significantly promoted venous thrombus resolution. Mechanistically, macrophages played crucial roles in venous thrombus resolution enhanced by caffeic acid via increasing macrophage recruitment and inhibiting macrophage M1 polarization in the thrombus. We also found that the beneficial effects of caffeic acid in the thrombus resolution involved in anti-inflammation effects, facilitating collagenolysis and neovascularization in the thrombus. Furthermore, we demonstrated that caffeic acid suppressed Keap1, thereby activating Nrf2 and promoting its nuclear translocation in macrophages to enhance venous thrombus resolution.

Previous studies have shown that caffeic acid has a high potential for treating and preventing cardiovascular diseases in preclinical studies, such as lowering hypertension, preventing atherosclerosis and myocardial infarction.^36^ A previous study has reported that caffeic acid inhibited photochemical injury-induced thrombosis in mouse cerebral arteries and ADP topical application-induced cerebral venular thrombosis *in vivo*, and alos inhibited ADP-induced platelet aggregation, P-selectin expression, ATP release, Ca^2+^ mobilization and integrin αIIbβ3 activation *in vitro*.^23^ Another study also found that caffeic acid significantly inhibited thrombin-induced platelet aggregation and fibrinogen binding to integrin αIIbβ3, and repressed platelet-mediated clot contraction *in vitro*.^37^ Taken together, these two studies indicated that caffeic acid could inhibit thrombus formation *in vivo* primarily through suppressing platelet function. Whereas, we found that caffeic acid promoted DVT resolution. One reason may be due to the different experimental murine models of thrombosis, The stasis-type DVT model involving surgical complete ligation of the mouse inferior vena cava was used in our study, while the mouse model of acute platelet mediated thrombosis including cerebral arteries thrombosis model induced by photochemical injury and cerebral venular thrombosis model induced by ADP topical application were used in the previous study, The other reason may be due to differences in drug dosage, with 40 mg/kg used in our study, 1.25-5 mg/kg used in the previous study.

Venous thrombus resolution is a critically important multistep process involving fibrinolysis, collagenolysis, phagocytosis of thrombus fragments by inflammatory macrophages (tissue clearing), neovasculation, and vessel wall remodeling.^6,38^ Macrophages are multifunctional leukocytes concerned with venous thrombus resolution.^8^ It was demonstrated that the increase of macrophage numbers or monocyte recruitment into the thrombus could improve VT resolution and recanalization.^30^ The number of macrophages was shown to start increasing and peaking at 7 days after IVC ligation and then gradually decreasing until the experimental deadline of 21 days.^39^ It was showed that caffeic acid inhibited LPS-induced M1 polarization of RAW264.7 cells *in vitro*.^24,40^ Consistently, our results showed that macrophage increased in the thrombus at 7 days after surgery and caffeic acid promoted macrophages recruitment into thrombus and inhibited macrophage M1 polarization, thereby promoting thrombus resolution. Furthermore, macrophages were confirmed to play a major role in the benefical effect of caffeic acid in thrombus resolution via macrophages depleted by chlorophosphate liposomes. Moreover, current studies has found that neutrophil extracellular traps (NETs) released by neutrophils are closely associated with DVT.^41^ Neutrophils are among the first immune cells to infiltrate the thrombus and set the stage for later thrombus resolution.^6^ Neutrophils predominantly present 1 day after IVC ligation,^31^ initially outnumbering monocyte/macrophage cells by a ratio of 7:1 at 24 hours and then steadily decreasing in number by approximately 50% per week.^39^ It was showed that caffeic acid modulated inflammatory activation of neutrophils and attenuated sepsis-induced organ injury.^42^ Considering that neutrophils enter the thrombus rapidly at 1-2 days contributing to early thrombus formation, while our study mainly focused on the 7^th^ day after IVC ligation when caffeic acid promoted thrombus resolution, thus whether caffeic acid affected the number of neutrophils infiltrating into the thrombus and the formation of NETs deserved further exploration.

Collagenolysis is a key event in the inflammatory vascular remodeling processes associated with venous thrombus resolution.^43^ In the present study, we found that caffeic acid promoted collagenolysis through increasing MMP-2 activity in the thrombus. Interestingly, caffeic acid was reported to repress MMP-2 and MMP-9 expression which was associated with obstruction of NF-κB activation in hepatocellular carcinoma, resulting in a reduction in cancer invasiveness and growth.^44^ It is possible that the discrepancy may be the result of the use of tumor cells. It was demonstrated that intrathrombotic macrophages could engulf necrotic tissue, clear cellular debris, and release proteolytic enzymes such as MMP-2, MMP-9, urokinase-type plasminogen activator (uPA), and tissue-type plasminogen activator (tPA), to dissolve the surrounding matrix and fibrin, thereby promoting thrombolysis and recanalization.^29,45^ However, uPA and tPA in the thrombus was not determined in our study. Therefore, whether caffeic acid has an effect on uPA and tPA fibrinolysis system to promote thrombus resolution needs further study.

Thrombus neovascularization has been demonstrated to be another key event during thrombus resolution and recanalization.^32^ Intrathrombotic macrophages express and release proangiogenic factors, such as VEGF, basic fibroblast growth factor, and placental growth factor, which stimulate capillary formation and neovascularization.^46,47^ It was shown that caffeic acid increases mRNA levels of VEGF in the brains of mice exposed to hypoxia.^48^ Our results also showed that caffeic acid increased the expression of VEGF and CD31, indicating that caffeic acid facilitated neovascularization in the thrombus. Notably, a previous study reported that caffeic acid suppressed hypoxia-induced VEGF mRNA expression and secretion in Caki-1 renal carcinoma cells and inhibited tumor growth and angiogenesis by inhibiting the expression of hypoxia inducible factor-1α and VEGF in mice bearing a Caki-1 carcinoma.^21^ We speculated that the opposite effect of caffeic acid on angiogenesis might be attributed to the use of tumor cell lines. However, how caffeic acid enhances neovascularization to promote thrombus resolution needs further research.

Numerous studies have shown that both inflammation and oxidative stress were central to throumbus resolution.^12,49^ Caffeic acid has been reported to display multiple biological activity, such as antioxidant, anti-inflammatory, immunomodulatory, etc.^50^ It was showed that caffeic acid alleviated metabolic-associated steatotic liver disease by reducing hepatic lipid accumulation, lipotoxicity and oxidative damage through activating Nrf2 via binding to Keap1.^51^ Moreover, it was reported that caffeic acid prevented acetaminophen-induced hepatotoxicity by decreasing Keap1 expression, inhibiting binding of Keap1 to Nrf2, and thus activating Nrf2 and leading to increased expression of antioxidative signals including heme oxygenase 1 and quinone oxidoreductase 1.^52^ Consistent with these previous studies, our RNA sequencing results showed that the expression of anti-oxidative stress-related genes was upregulated and while inflammation-related genes were down-regulated in BMDMs treated with caffeic acid. Moreover, our above results showed that caffeic acid inhibited macrophage M1 polarization *in vivo* and *in vitro*, also confirming the anti-inflammatory activity of caffeic acid. Furthermore, we found that caffeic acid enhanced thrombus resolution through targeting the Keap1-Nrf2 axis: Specifically, it inhibited Keap1-dependent cytoplasmic retention of Nrf2, enabling Nrf2 nuclear translocation and subsequent activation of downstream antioxidant pathways in macrophages. However, further investigation is needed to elucidate how caffeic acid modulates Keap1-the key negative regulator of Nrf2-to facilitate its nuclear translocation during thrombus resolution. Future studies should explore whether caffeic acid directly binds to Keap1 to disrupt its interaction with Nrf2 or if it affects upstream signals that regulate Keap1’s inhibitory activity. Clarifying these molecular interactions could uncover novel therapeutic targets to enhance thrombus resolution.

In conclusion, our study demonstrated that caffeic acid promoted venous thrombus resolution by inhibiting Keap1 and activating Nrf2, mainly through facilitating collagenolysis as well as neovascularization in thrombus. Our findings also emphasized on the vital role of macrophages in caffeic acid enhancing venous thrombus resolution. Our study provides a new evidence to highlight the therapeutic potential of caffeic acid for the treatment of deep vein thrombosis.

## Acknowledgments

This study was supported by grants from the National Natural Science Foundation of China (No 82204737), the Science and Technology Commission of Shanghai Municipality (No 24ZR1465300), the Health Commission of Shanghai Municipality (ZY(2021-2023)-0203-04), Future Plan for Traditional Chinese Medicine Inheritance and Development of Shanghai Municipal Hospital of Traditional Chinese Medicine (WLJH2021ZY-ZYY007; WL-HBBD-2021001K), Traditional Chinese Medicine Research Project of Hongkou District Health Commission (HKZYY-2024-06), Medical Research Project of Hongkou District Health Commission (HongWei 2402-03)

## Author contribution

Zhou-yu NIE, Jia-qi ZHANG, and Jia-yi SHEN-YUAN contributed to the conception and design of the study, and performed the research; Meng-Jiao LU, Yang SHEN and Liang SHI contributed to the acquisition, analysis and interpretation of data; Zhou-yu NIE drafted the manuscript. Ling LI, Li-chao ZHANG and Yong-Bing CAO supervised the project and revised the manuscript critically for important intellectual content. Professor Li-li Ji from Shanghai University of Traditional Chinese Medicine provided Nrf2 gene-deficient mice. All authors have approved the final vision of this manuscript.

## Abbreviations

CA: caffeic acid
DVT: Deep vein thrombosis
MMP-2 and 9: matrix metalloproteinases-2 and 9
BMDM: bone marrow-derived macrophages
Nrf2: nuclear factor erythroid 2-related factor 2
Keap1: Kelch-like ECH-associated protein 1
CD68: cluster of differentiation 68
iNOS: inducible nitric oxide synthase
Arg-1: Arginine-1
H&E: hematoxylin and eosin
GO: Gene Ontology
KEGG: Kyoto Encyclopedia of Genes and Genomes
LPS: lipopolysaccharide
IFN-γ: interferon-γ
IL-1β: interleukin-1β
IL-4: interleukin-4
IL-13: interleukin-13
TGF-β: transforming growth factor-β
CD31: Platelet endothelial cell adhesion molecule-1
VEGFA: vascular endothelial growth factor A

---

**Supplementary Figure 1.**
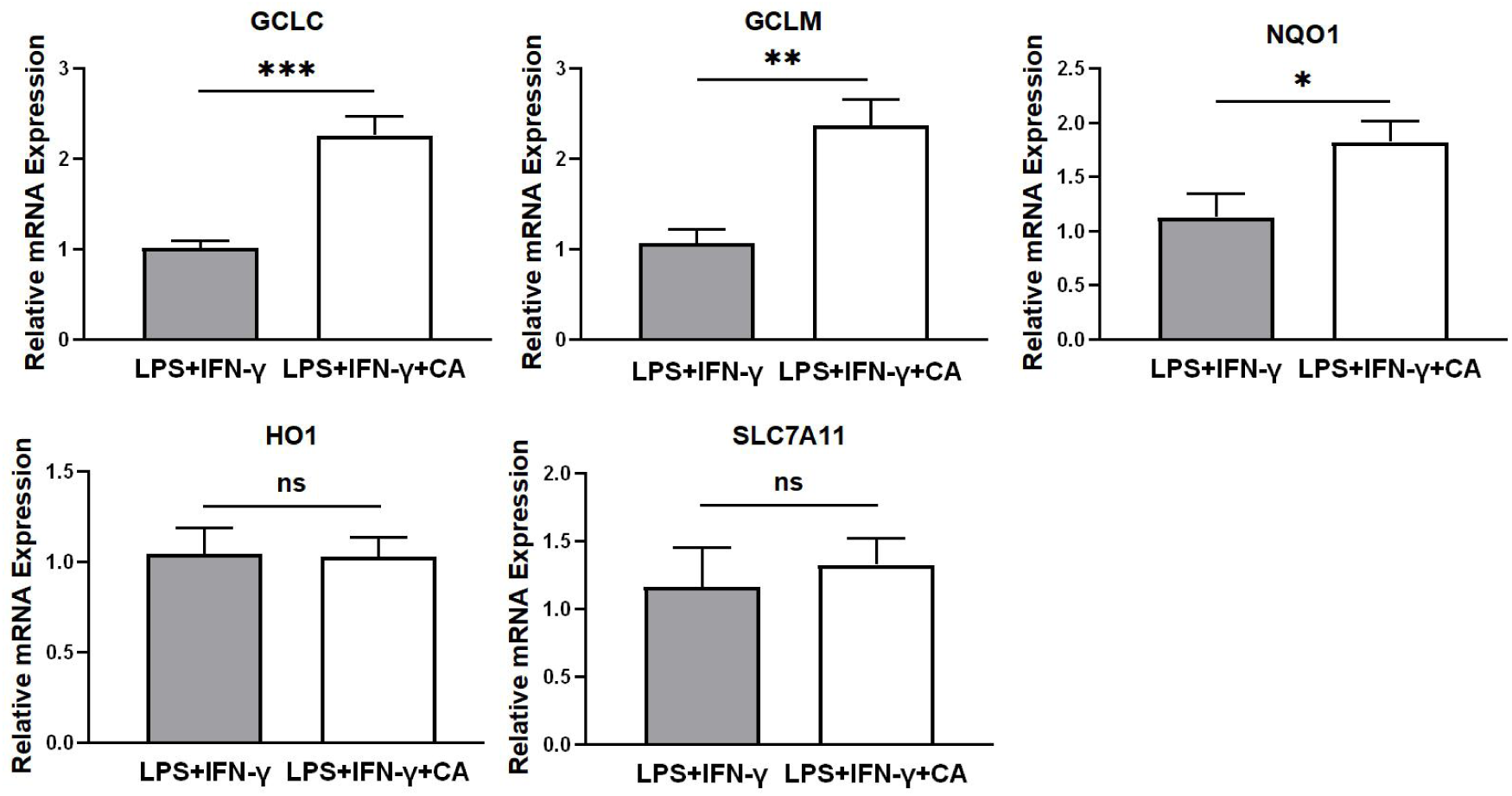
Representative qRT-PCR for Gclc, Gclm, Nqo1, Ho1, Slc7a11 expression in bone marrow-derived macrophages induced by LPS and IFN-γ for 24 hours in vitro.

